# Rapid estimation of photon measurement density functions using a deep convolutional neural network for functional near-infrared spectroscopy

**DOI:** 10.64898/2026.04.12.717947

**Authors:** Yang Zhao, Xue-Ting Sun, Wen-Di Shi, Chao-Zhe Zhu, Lei Zhang

**Author notes:** Address correspondence to Lei Zhang, Ph.D. Chinese Institute for Brain Research, Beijing (CIBR), Bldg.3, NO.9, YIKE Road, Zhongguancun Life Science Park, Changping District, Beijing, China.

## Abstract

The photon measurement density function (PMDF) plays a fundamental role in both pre-experimental optode arrangement and post-experimental data analysis in functional near-infrared spectroscopy (fNIRS). Conventionally, PMDFs are derived from structural MR images through tissue segmentation and photon propagation modeling (PPM), which are computationally demanding and time-consuming, thereby limiting their practical use. In this study, we propose a novel deep learning–based framework to estimate PMDFs directly from MR images and channel configurations. The proposed method supports flexible source–detector distances and eliminates the need for explicit tissue segmentation and repeated photon simulations. Specifically, a convolutional neural network is trained to predict photon fluence distributions, from which PMDFs are subsequently derived using the adjoint formulation. The trained model is evaluated on channels placed in both trained and unseen scalp regions across commonly used source–detector distances. The results demonstrate that the proposed method achieves PMDF estimations comparable to those obtained from PPM. Overall, this approach significantly reduces computational cost and has the potential to facilitate broader adoption of PMDF-based methods in the fNIRS community.

## 1. Introduction

Functional near-infrared spectroscopy (fNIRS) is a non-invasive optical neuroimaging technique which has been widely applied in both scientific research and clinical settings in recent years (Rahman et al., 2020; Scholkmann et al., 2014). Compared to other neuroimaging modalities, fNIRS offers advantages such as high ecological validity, low sensitive to head motion, portability, and cost efficiency (Pinti et al., 2020). These features make it particularly suitable for studies involving special populations including infants and elderly, as well as studies conducted in naturalistic environments (Gervain et al., 2023; Mei et al., 2023; Zhao et al., 2024).

In fNIRS, measurements are acquired through channels formed by a source–detector pair, with a typical source–detector distance up to approximately 45 mm (Frijia et al., 2021). The recorded signals depend on the propagation of near-infrared light through biological tissue, commonly illustrated as a “banana-shaped” photon path. More formally, this photon path is described by the photon measurement density function (PMDF), a spatial function that characterizes the sensitivity of the detector to optical property changes at each location within the tissue underlying a given channel (Arridge, 1995; Arridge and Schweiger, 1995).

The PMDF has been increasingly adopted in fNIRS research in recent years, as it provides a more realistic representation of light propagation compared with simplified approaches such as the point-to-point model, which projects the midpoint between source and detector to cortical surface (Okamoto and Dan, 2005). While the point-based model offers a computationally efficient approximation, it does not fully capture the spatially distributed nature of photon transport in biological tissue (Brigadoi et al., 2018; Cai et al., 2021). In experimental design, the PMDF has been used to optimize optode arrangement, either based on individual structural MRI data or standardized anatomical templates such as MNI and Colin27 (Brigadoi et al., 2018; Machado et al., 2018; Morais et al., 2018). In data analysis, PMDF-based approaches have been applied to investigate the relationship between fNIRS and fMRI signals and to improve the accuracy and reliability of activation mapping (Duan et al., 2012; Sassaroli et al., 2006; Zhai et al., 2020). Furthermore, in diffuse optical tomography (DOT), the PMDF is essential for constructing the forward model and solving the associated inverse problem (Vidal-Rosas et al., 2023).

The conventional pipeline for estimating the PMDF based on a structural MRI typically involves two main steps: head tissue segmentation, followed by photon propagation modelling (PPM) using numerical methods such as the Monte Carlo simulation or the finite element method (FEM) for solving the diffusion equation (Arridge et al., 1993; Boas et al., 2002). These procedures are computationally intensive and time-consuming. Accurate PMDF estimation depends on high-quality tissue segmentation, which often requires sophisticated algorithms and may take several hours to complete. Monte Carlo simulations usually require a large number of photons (typically >10^7^) to achieve sufficient accuracy, leading to substantial computational cost. Although GPU-based parallel computing can accelerate this process, the overall burden remains considerable (Fang and Boas, 2009; Fernandes et al., 2026). Therefore, the computational complexity of PMDF estimation limits its broader application in the current fNIRS community.

This limitation becomes particularly critical in applications such as optode arrangement optimization. To effectively probe regions of interest (ROIs), fNIRS optodes can be flexibly placed at arbitrary scalp locations, resulting in a large number of possible channel configurations. Each configuration requires an associated PMDF to evaluate its sensitivity to ROIs, thereby necessitating repeated photon propagation simulations. As a result, PMDFs must be computed for a large number of channels, significantly increasing the computational demand. Moreover, recent studies suggest that PMDFs derived from multiple subjects provide better generalization to individuals without MRI data than those based on a single template, further multiplying the number of required simulations (Zhao et al., 2025). Therefore, a computationally efficient and scalable approach for PMDF estimation is highly desirable for advancing fNIRS research and applications.

Deep learning has emerged as a powerful tool in neuroimaging studies, especially for learning complex mappings from anatomical images to underlying physical or physiological fields (Litjens et al., 2017; Shen et al., 2016). It has been successfully applied in fNIRS studies, including brain–computer interfaces and the clinical diagnosis of neuropsychiatric disorders (Eastmond et al., 2022). Notably, recent work has demonstrated the use of deep convolutional neural networks (CNNs) to predict electric fields induced by transcranial magnetic stimulation (TMS), suggesting the potential of data-driven approaches for modeling complex physical processes in the brain (Yokota et al., 2019). Inspired by these advances, we propose a deep learning–based approach for efficient PMDF estimation in fNIRS. The proposed method directly estimates PMDFs from structural MR images and channel configurations, without requiring tissue segmentation and photon propagation modeling. Moreover, it supports varying source–detector distances, enabling flexible application across different channel configurations. We validate the proposed approach at two commonly studied scalp areas, and the results demonstrate that it can generate PMDFs comparable to those obtained using conventional numerical methods, while substantially reducing computational cost.

## 2. Materials and Methods

### 2.1. General workflow of CNN-based PMDF estimation

The goal of the proposed method is to estimate the PMDF given a subject-specific MR image and a channel configuration, defined by the source and detector locations on the scalp. This task can be formulated as a supervised learning problem. A straightforward approach is to train a neural network model to directly predict the PMDF from the MR image and channel configuration. However, since channels can be placed at arbitrary scalp locations with varying source–detector distances, this formulation leads to high input dimensionality and increased model complexity. An alternative strategy is that instead of directly predicting the PMDF given a channel configuration, we first train a model to estimate the photon fluence distribution given an MR image and a single optode location. The PMDF can then be derived using the adjoint formulation based on the predicted fluence distributions of the source and detector (Arridge and Schweiger, 1995). This design reduces the dimensionality of the problem, simplifies the model, and enables flexible channel configurations.

Specifically, in the training stage, a neural network model *f* is trained to predict the photon fluence Φ from the input MR image *x* and optode location *s*, where (*x, s*, Φ) ∈ 𝒟_*train*_ (Fig. 1A). The ground-truth fluence distributions are obtained using photon propagation modeling. In the testing stage, the PMDF estimation consists of two steps. First, given an unseen MR image *x* and a channel defined by source and detector locations *s* and *d*, i.e., (*x, s, d*) ∈ 𝒟_*test*_, the corresponding photon fluence distributions of source 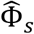 and detector 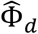 are predicted by the trained neural network model 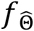 with the estimated parameters 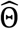 (Fig. 1B). Second, the PMDF is computed using the adjoint formulation:

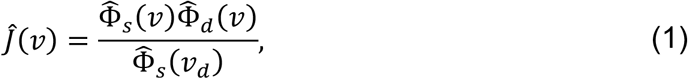

where 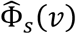 and 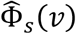 denote the predicted photon fluence at voxel *v* generated by the source and detector, respectively. 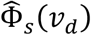 represents the photon fluence at the detector location *v*_*d*_ produced by the source and serves as the normalization factor. *Ĵ*.(*v*) denotes the estimated PMDF at voxel *v*.

**Figure 1.**
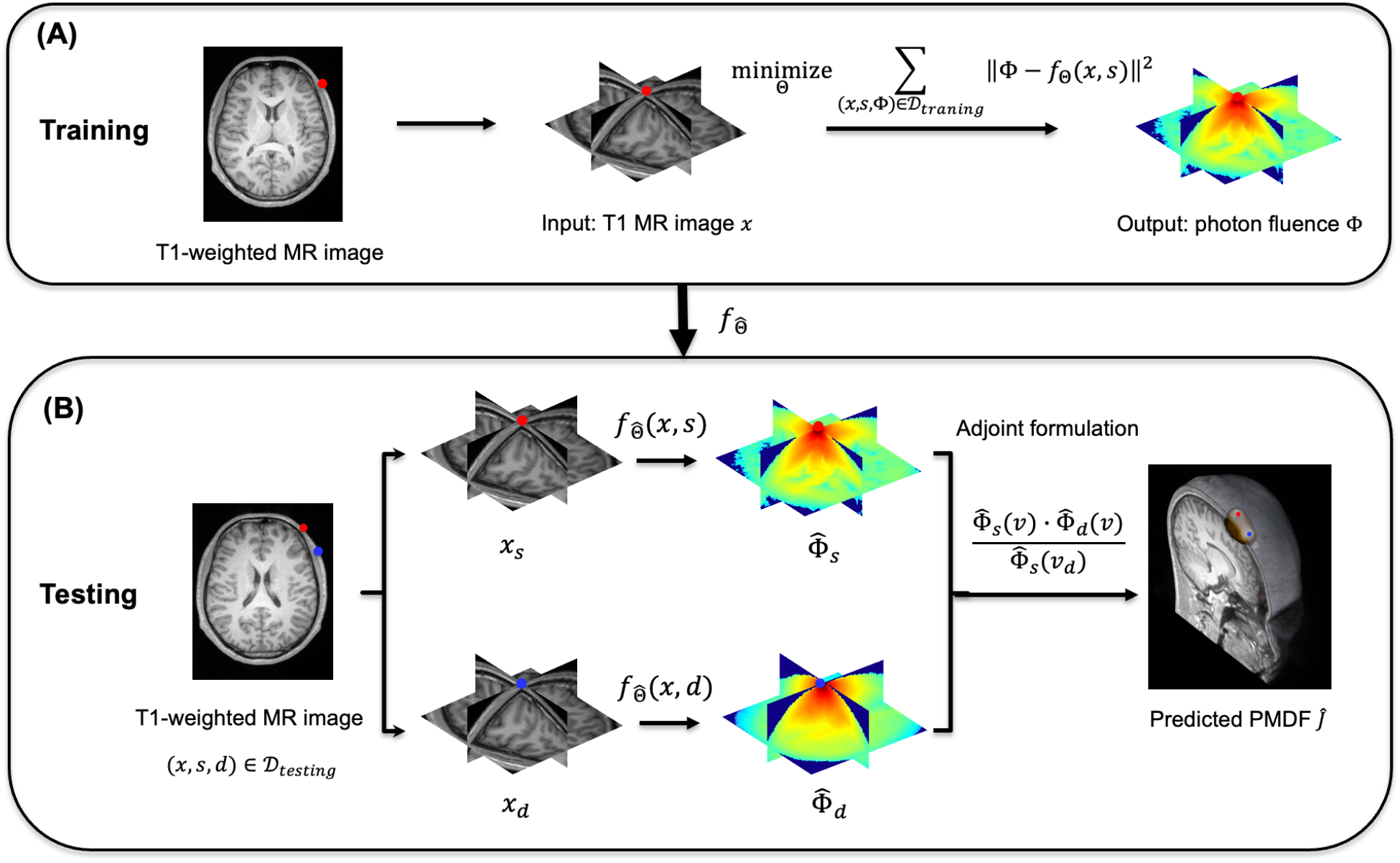
Workflow of the CNN-based PMDF estimation framework using subject-specific MR images. (A) Training stage: For each scalp location across subjects in the training dataset, cuboidal subvolumes of MR images and corresponding photon fluence distributions are extracted based on the optode-centered coordinates. A convolutional neural network (CNN) is trained to learn the nonlinear mapping from MR image subvolumes to photon fluence distributions. (B) Testing stage: For a given channel placement on an unseen subject, the subvolumes of source (Red dot) and detector (Blue dot) are extracted based on their scalp locations. The corresponding photon fluence distributions are predicted using the trained model. The PMDF is then computed using the adjoint formulation based on the estimated source and detector fluence.

### 2.2. MRI and PPM dataset

We utilized a previously published dataset comprising structural MR images of healthy subjects and corresponding photon propagation modelling results (Wei et al., 2025). Specifically, the dataset includes 47 T1-weighted MR images (23 male 24 female). The MR images were segmented into five tissue types — scalp, skull, cerebrospinal fluid (CSF), gray matter and white matter, and subsequently reconstructed into tetrahedral meshes using the SIMNIBS software (Thielscher et al., 2015). A uniformed scalp sampling method was employed to generate high-resolved optode locations (approximately 18,000 per subject) based on the continuous proportional coordinate (CPC) system (Xiao et al., 2018). These sampling points uniformly cover the scalp with an average inter-point distance of approximately 2 mm. For each sampled scalp locations and each subject, the photon propagation modelling was performed using mesh-based Monte Carlo simulation (MMC) software (Fang, 2010), resulting in approximately 18000 photon fluence distributions per subject. Since we estimate PMDFs in voxel space in this study, the simulated photon fluence distributions were mapped from the mesh domain to the voxel space using a point-in-tetrahedron method. The optical properties of each tissue type were assigned based on averaged values across commonly used near-infrared wavelengths (Strangman et al., 2014). Detailed modeling parameters can be found in the previously published study (Wei et al., 2025).

### 2.3. Model training

To train the model for predicting photon fluence distributions from MR images, data from *N*_*x*_ *=* 37 out of 47 subjects were randomly selected as the training set. A subset of scalp locations was defined by selecting points within a circular region centered at C3 (according to the international 10–20 system) with a radius of 30 mm on a representative subject (Fig. 2A). This region was selected because it overlies the motor cortex, which has been widely investigated in neuroimaging studies. This resulted in 637 scalp locations, which were then mapped to other subjects using the corresponding CPCs. Due to the exponential decay of photon fluence with increasing distance from the source, the fluence distributions are typically sparse. To reduce computational cost and improve training efficiency, subvolumes are extracted from both the MR images and the corresponding photon fluence distributions (Fig. 2C, D). The extraction is defined based on optode locations and orientations using the Scalp Geometry-based Parameter (SGP) coordinate system (Jiang et al., 2022). Since PMDFs are derived using the adjoint formulation, which involves the product of source and detector photon fluence distributions, only overlapped voxels between the two subvolumes can be estimated. Therefore, the subvolume size must be sufficiently large to allow predictions of commonly used source–detector distances (up to 45 mm). In this study, a subvolume size of 104 × 104 × 48 *mm*^3^ was adopted. We further evaluated whether this subvolume size is sufficient to capture the relevant PMDF regions for typical channel distances. Details of the cuboid size validation are provided in the Supplementary Materials (Section 6.1). The results indicate that it adequately supports PMDF estimation for source–detector distances up to 45 mm. To augment the training data, 36 orientations were generated for each scalp location, resulting in *N*_*s*_ *=* 637 × 36 samples per subject (Fig. 2B). Consequently, the total number of training samples is *N = N*_*x*_ × *N*_*s*_ *=* 848,484. Finally, due to the high dynamic range of photon fluence distributions, a logarithmic transformation is applied prior to training:

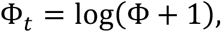

to stabilize the training process (Ardakani et al., 2022).

**Figure 2.**
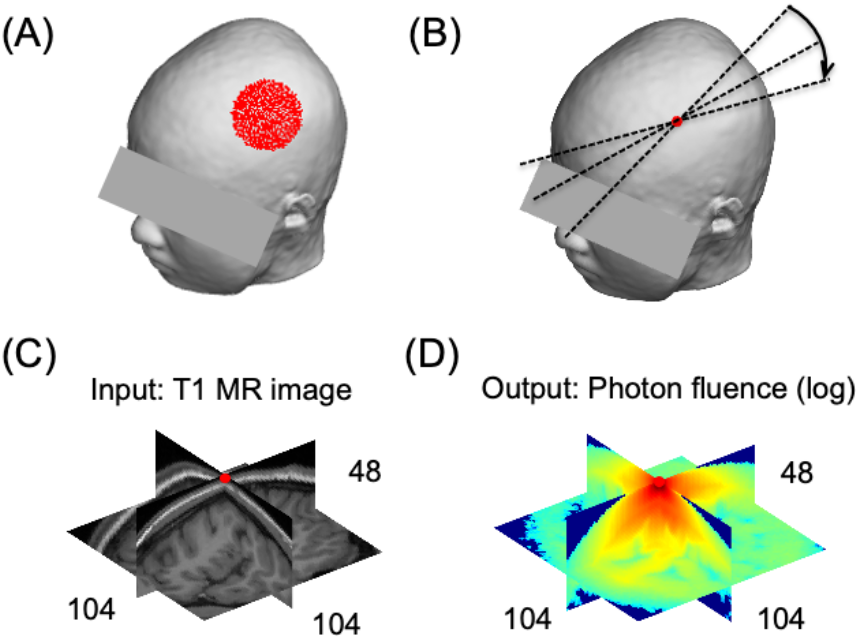
Construction of the training dataset. (A) Scalp sampling locations (red dots) on an example subject used for training. (B) Data augmentation through rotation of optode orientations at a typical scalp location. (C) Extracted MR image subvolume of size 104×104×48 mm^3^. (D) Corresponding photon fluence subvolume extracted using the same optode-centered coordinate system.

The training process can be formulated as a regression problem, in which the discrepancy between the model output and the ground-truth photon fluence is minimized by optimizing the model parameters. Specifically, the optimization problem can be written as:

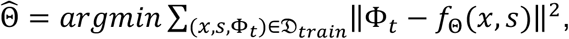

where Φ _*t*_ denotes the ground-truth photon fluence, and *f*_Θ_ (*x, s*) represents the model output given an MR image subvolume *x* and an optode location *s* with parameters Θ. The training samples consist of paired MR image subvolumes and their corresponding photon fluence distributions for different scalp locations in the training dataset.

Convolutional neural networks (CNNs) provide a powerful framework for solving such regression problems and have demonstrated strong performance in various medical imaging applications (Chen et al., 2025). Among these, the U-Net architecture has been widely adopted, particularly for prediction electric field induced by TMS (Yokota et al., 2019). U-Net consists of a contracting path that captures contextual information through successive convolution and pooling operations, and an expanding path that enables precise spatial localization via upsampling and convolution. The skip connections allow the network to integrate high-level semantic features with low-level spatial details. In this study, A 3D U-Net architecture with a depth of 4 was employed (Fig. 3). The network follows an encoder–decoder structure with skip connections, using repeated convolutional blocks, batch normaliztion, and nonlinear activation functions. The model was trained for 20 epochs with a batch size of 20, using stochastic gradient descent (SGD) as the optimizer and mean squared error (MSE) as the loss function.

**Figure 3.**
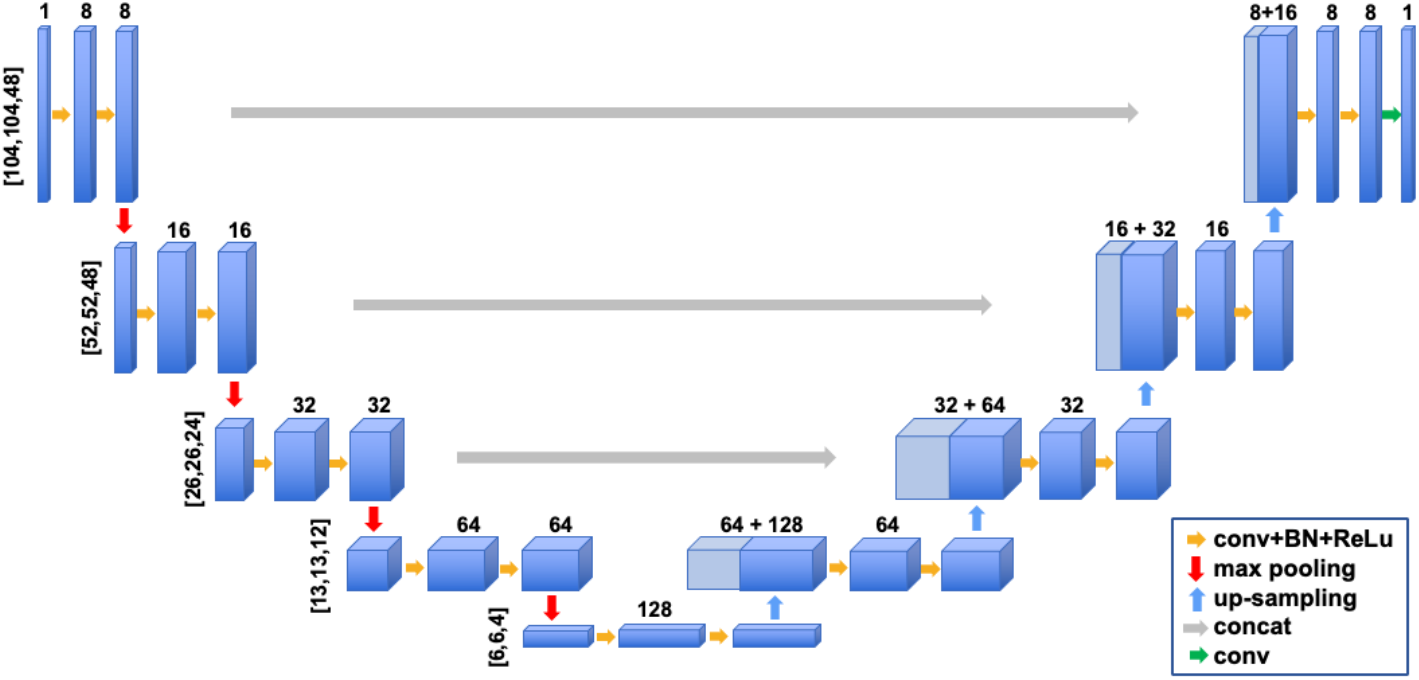
Architecture of the convolutional neural network (CNN), specifically a U-Net with a depth of 4, used for photon fluence estimation.

Based on the trained model, the PMDF can be estimated for an unseen subject given an MR image *x* ∈ 𝒟_*test*_ and channel configuration defined by source and detector locations *s* and *d*, where *s, d* ∈ 𝒟_*test*_. The MR image is first cropped into subvolumes centered at the source and detector positions using the corresponding optode-centered coordinate system. The trained model *f* with optimized parameters 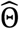 is then used to predict the logarithm-transformed photon fluence distributions for both source and detector locations, i.e.,

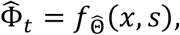

and similarly for the detector location *d*. The predicted photon fluence is subsequently recovered via inverse transformation:

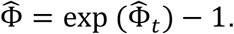

Since the predicted photon fluence is defined in the optode-centered coordinate system, it is resliced back to the original MR image space. Finally, the PMDF is computed using the adjoint formulation (Eq. 1).

### 2.4. Performance evaluation

To evaluate the performance of the proposed framework, the remaining 10 subjects were used as the test set, with their MR images and corresponding PMDFs serving as ground truth. The ground-truth PMDFs were derived from photon fluence distributions obtained via photon propagation modeling (PPM) using Eq. (1). Since fNIRS signals primarily originate from cerebral tissues, only PMDF values within the gray matter were included in the evaluation.

We first evaluated PMDFs for channels with a commonly used source–detector distances of 30 mm at the predefined scalp locations. The channel centers were placed at the sampled scalp locations in the motor cortex region, with 12 orientations per location. To ensure that channels remained within the training region, configurations with optodes located outside the sampled scalp area were excluded, resulting in approximately 3,000 channels per subject.

The prediction accuracy was assessed at multiple levels. At the voxel level, the mean absolute error (MAE) of the predicted channel PMDF was defined as:

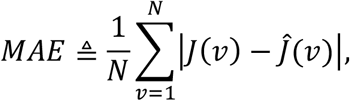

where *J*(*v*) and Ĵ.(*v*) denote the ground-truth and predicted PMDF values at voxel *v*, respectively, and *N* is the number of voxels considered. In addition, the mean relative error (MRE) was defined as:

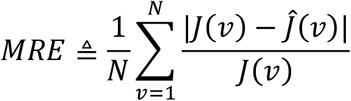

Since PMDF values in intracerebral regions are typically small (e.g., < 0.1), including all voxels would lead to artificially low MAE values and high MRE. Therefore, only voxels whose ground truth value exceeding a threshold were included in the voxel-level evaluation.

In addition to voxel-level analysis, ROI-level evaluation was performed. Brain regions were defined based on a standard atlas, the Brodmann atlas in the MNI space (Pijnenburg et al., 2021). The Brodmann areas were obtained on individual MR image of each subjects based on the automatic anatomical registration. The sensitivity to each ROI was computed by summing PMDF values within the Brodmann areas. As a given channel may be sensitive to multiple areas, the ROI with the maximum sensitivity was selected for evaluation.

To assess the generalizability of the proposed method, we further evaluated PMDF prediction for channels placed in a different scalp region. Specifically, channels at the frontal region were selected using F3 (according to the 10–20 system) as the center and the same configuration as the C3-centered setup.

To evaluate performance across different channel configurations, the proposed method was tested on a range of source–detector distances from 15 mm to 45 mm, with a step size of 5 mm. Distances below 15 mm were excluded, as such short distances are generally considered to have limited sensitivity to brain cortex.

Analytical models have also been widely adopted for estimating PMDFs in fNIRS studies, as they avoid the need for computationally intensive simulations (Boas et al., 2002; Feng et al., 1995). In these approaches, the head is typically modeled as a semi-infinite homogeneous medium, and photon propagation is described using diffusion theory. Under this assumption, the photon fluence generated by a source located at r_*s*_ can be expressed as:

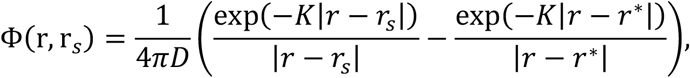

where *D* is the diffusion coefficient, *K* is the effective attenuation coefficient, *r* denotes a position within the medium, and *r*^*\**^ represents the location of the image source introduced to satisfy boundary conditions. To compare the proposed method with the analytical model, the above formulation was used to compute photon fluence, and the PMDF was subsequently derived using the same adjoint formulation as in Eq. (1). The optical parameters *D* and *K* were determined based on the absorption coefficient *μ*_*s*_ *=* 0.11 *mm*^97^ and reduced scattering coefficient *μ*_*a*_ *=* 0.2 *mm*^97^, following values reported in the literature (Sasai et al., 2012; Sassaroli et al., 2006).

## 3. Results

### 3.1. Voxel-level prediction accuracy

To evaluate whether the proposed CNN-based framework can accurately estimate PMDFs conditioned on subject-specific MR images, the predicted PMDFs were resliced and aligned in a channel-centered coordinate system (Wei et al., 2025). In this coordinate system, the origin is defined as the midpoint between the source and detector, the x-axis is aligned with the source–detector direction, and the z-axis is perpendicular to the scalp surface (Fig. 4). Figure 4A shows the voxel-wise correlation coefficients between PMDFs estimated by the CNN and PPM under this coordinate system. High correlation values are observed in regions beneath the channel, indicating that the proposed model can reliably capture spatial variations in PMDFs based on the underlying MR images. The depth-wise slices further demonstrate that high correlations persist in deeper regions (up to approximately 25 mm below the scalp), covering the cortical gray matter. Figure 4B provides a scatter plot comparing voxel-wise PMDF values from the two methods. The strong agreement between the predicted and ground-truth values further confirms the accuracy of the proposed approach. Notably, higher correlations are observed in regions near the center of the channel, where PMDF values are larger compared to peripheral areas. These regions are of primary importance in fNIRS, as they contribute most significantly to the measured signals.

**Figure 4.**
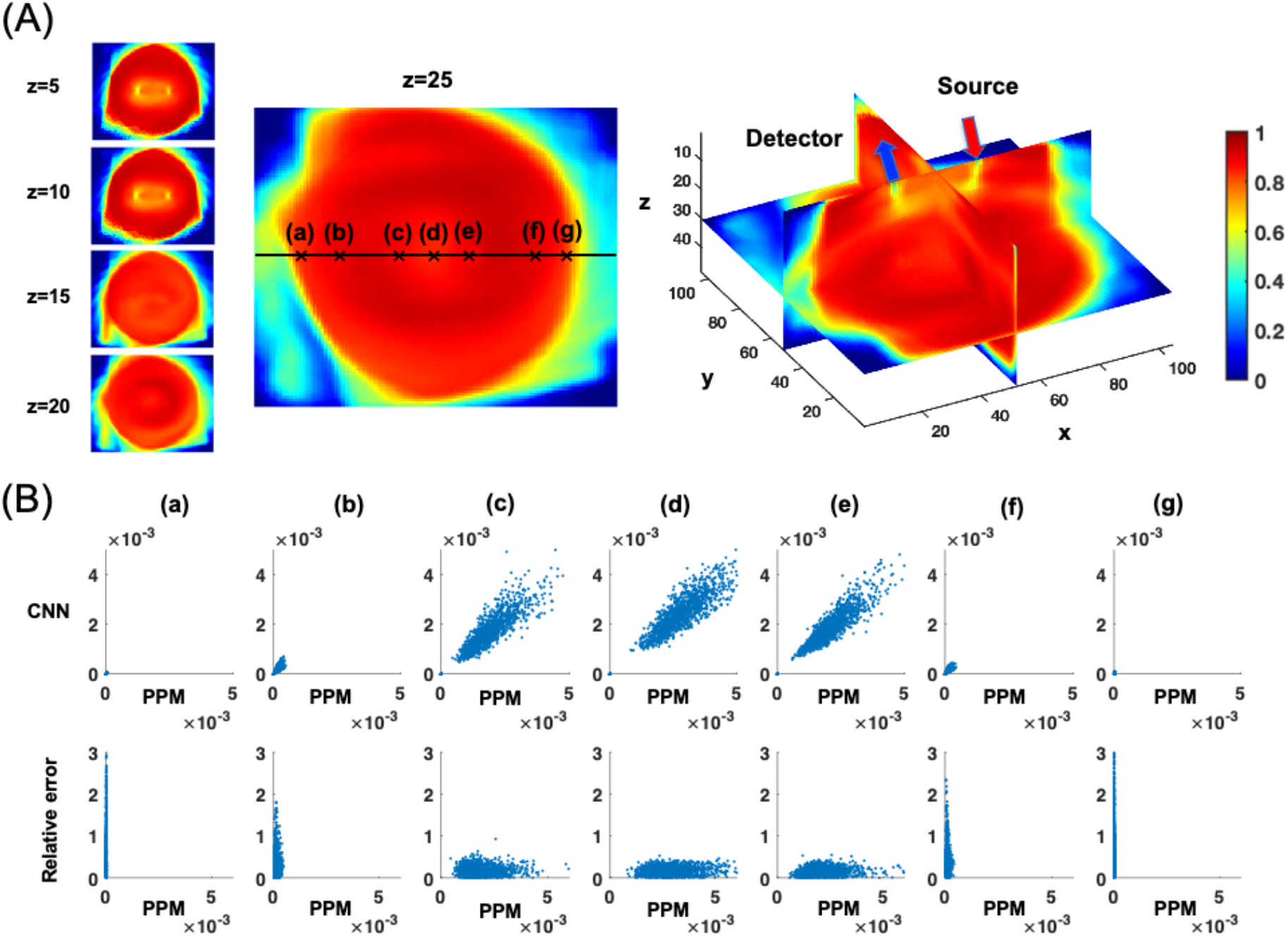
Voxel-wise correlation analysis of PMDFs estimated by photon propagation modeling (PPM) and the proposed CNN-based method in the motor area for a source–detector distance of 30 mm. (A) Spatial distribution of voxel-wise correlation coefficients in the channel-centered coordinate system. PMDFs estimated by both methods are aligned under the same coordinate system prior to correlation analysis. (B) Scatter plot comparing PMDF values estimated by CNN and PPM for voxels highlighted in the slice shown in (A).

To visualize the prediction performance on the cortical surface, the PMDFs predicted by the proposed framework were projected back onto the mesh space for six randomly selected subjects and source-detector distance of 30 mm (Fig. 5). Clear similarity is observed between PMDFs estimated by the CNN and PPM in the motor cortex (Fig. 5A). The high accordance is also confirmed by the small absolutely error between PMDFs derived by the two methods. By using a consistent color scale across subjects, the results also demonstrate that the proposed model is capable of accurately predicting not only the spatial patterns but also distinguishing the magnitudes of PMDFs across subjects. In the frontal cortex (Fig. 5B), a similar overall pattern is observed. However, slightly higher absolute errors are present compared to the motor cortex. This difference may be attributed to the fact that the model was primarily trained on data from the motor cortex region. The CNN-based method demonstrates high prediction accuracy, as indicated by low MAE and MRE values, both of which are substantially lower than those of the analytical model (Table 1). Note that the PMDF values included in the calculate the MAE and the MRE were both tresholded at 0.01. Other threshoulds were also tested and the results shows ignorance differences.

**Table 1.** Mean absolute error and relative mean absolute error of the CNN-based method and the analytical model at both voxel- and ROI-levels across multiple source-detector distances.

| Motor area |  |  |  |  |
| --- | --- | --- | --- | --- |
| SD (mm) | CNN |  | Analytical model |  |
| | Voxel-level<br>MAE ($\times 10^{-2}$ mm) /MRE | ROI-level<br>MAE (mm) /MRE | Voxel-level<br>MAE ($\times 10^{-2}$ mm) /MRE | ROI-level<br>MAE (mm) /MRE |
| 15 | 0.24/0.21 | 0.85/0.18 | 0.88/0.75 | 3.09/0.78 |
| 20 | 0.24/0.19 | 1.65/0.16 | 1.03/0.79 | 7.13/0.81 |
| 25 | 0.23/0.16 | 2.14/0.13 | 1.19/0.83 | 11.91/0.82 |
| 30 | 0.23/0.15 | 2.62/0.12 | 1.28/0.83 | 15.86/0.82 |
| 35 | 0.25/0.16 | 3.55/0.14 | 1.3/0.82 | 18.52/0.82 |
| 40 | 0.3/0.19 | 4.76/0.17 | 1.28/0.8 | 18.79/0.81 |
| 45 | 0.34/0.22 | 6.08/0.2 | 1.25/0.78 | 20.57/0.79 |
| Frontal area |  |  |  |  |
| SD (mm) | CNN |  | Analytical model |  |
| | Voxel-level<br>MAE ($\times 10^{-2}$ mm) /MRE | ROI-level<br>MAE (mm) /MRE | Voxel-level<br>MAE ($\times 10^{-2}$ mm) /MRE | ROI-level<br>MAE (mm) /MRE |
| 15 | 0.38/0.29 | 2.54/0.3 | 1/0.76 | 6.02/0.79 |
| 20 | 0.41/0.27 | 4.64/0.26 | 1.22/0.81 | 12.83/0.81 |
| 25 | 0.4/0.23 | 5.84/0.21 | 1.38/0.82 | 19.95/0.82 |
| 30 | 0.41/0.23 | 7.15/0.2 | 1.41/0.81 | 25.12/0.81 |
| 35 | 0.44/0.25 | 8.89/0.21 | 1.37/0.79 | 28.22/0.79 |
| 40 | 0.45/0.26 | 10.07/0.22 | 1.28/0.75 | 29.2/0.76 |
| 45 | 0.47/0.28 | 11.42/0.23 | 1.22/0.72 | 29.43/0.72 |

**Figure 5.**
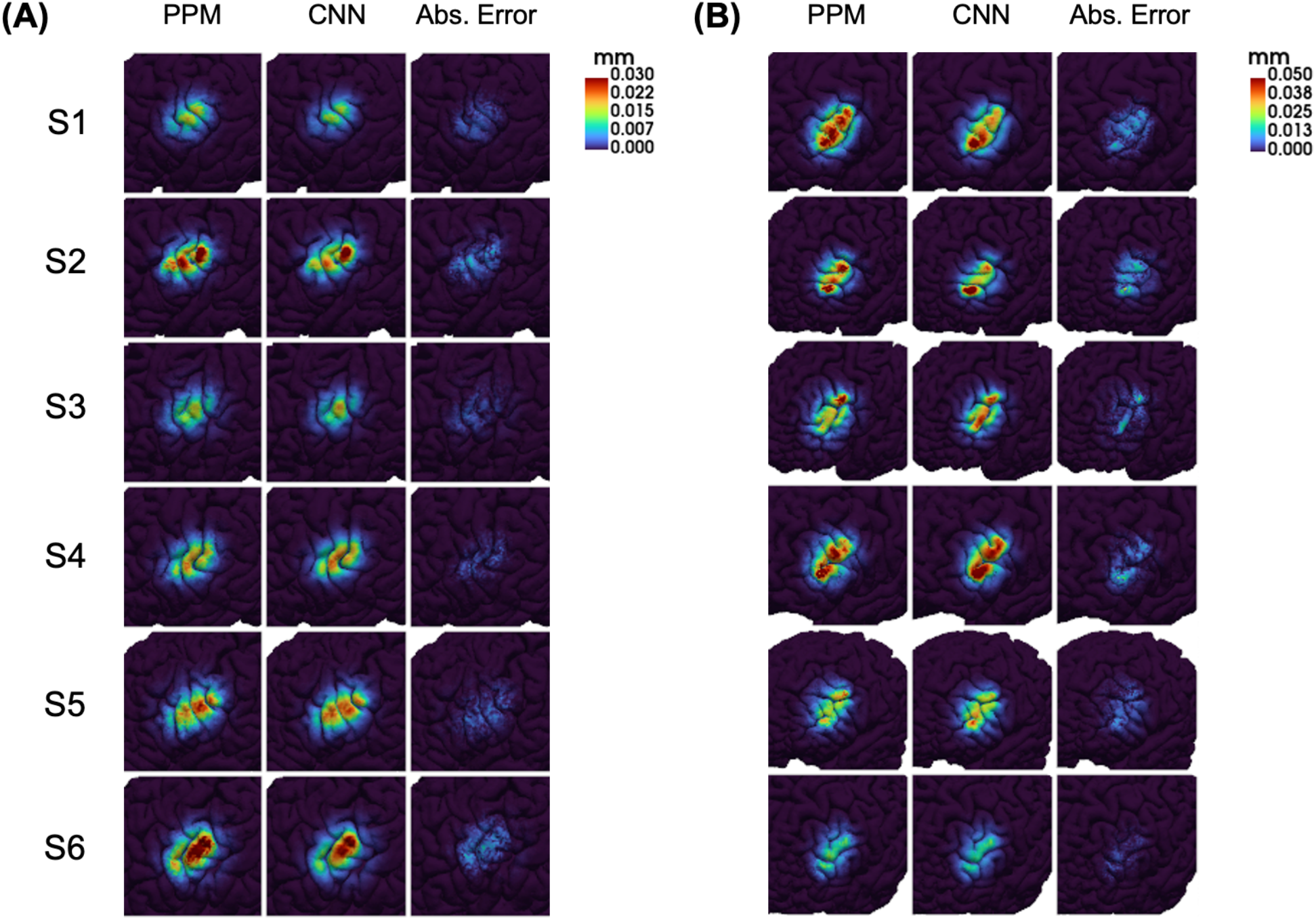
Comparison of PMDFs for channels with a source–detector distance of 30 mm estimated by the CNN and photon propagation modeling (PPM) on the cortical surface of six randomly selected subjects from the test dataset. (A) PMDF comparison in the motor cortex. (B) PMDF comparison in the frontal cortex.

### 3.2. ROI-level prediction accuracy

Figure 6 presents the ROI-level sensitivity of channels placed in the motor and frontal regions for a randomly selected subject. For each scalp location, the sensitivity is computed as the average sensitivity to the target ROI (highlighted in yellow) across all tested orientations. In both regions, the sensitivities predicted by the CNN-based method show strong agreement with those obtained from PPM in terms of both spatial distribution and magnitude. The scatter plots further demonstrate a high correlation between the two methods. In the frontal region, the scatter distribution appears slightly more dispersed but the overall agreement remains satisfactory. Consistent with the voxel-level evaluation, the proposed method achieves substantially lower mean absolute error (MAE) and relative MAE (RMAE) compared to the analytical model across different source–detector distances (Table 1). These results indicate that the CNN-based approach can accurately estimate ROI-level sensitivity, supporting its potential application in optode arrangement optimization.

**Figure 6.**
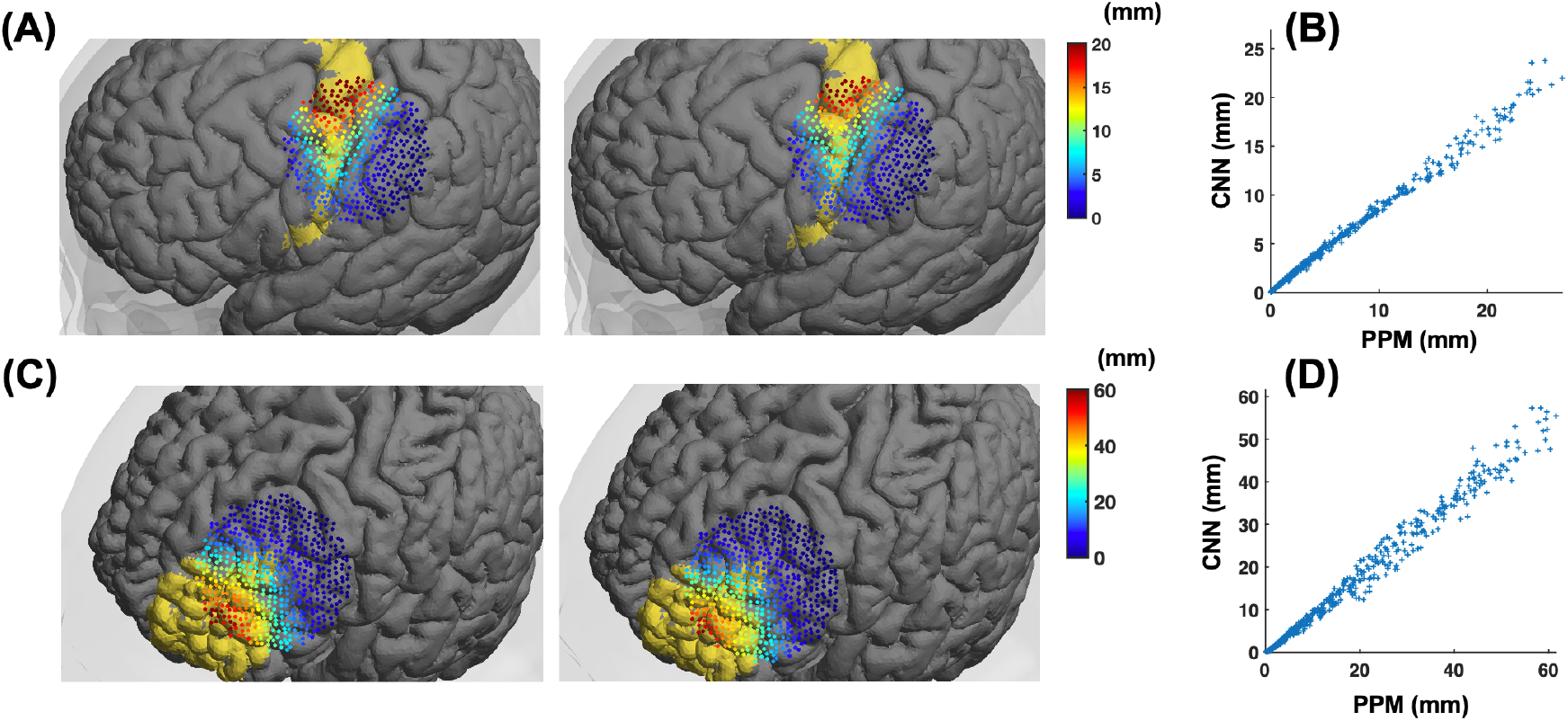
ROI-level sensitivity analysis for a randomly selected subject with a source–detector distance of 30 mm. Sensitivity to the primary motor cortex (Brodmann area 4, top row) and anterior prefrontal cortex (Brodmann area 10, bottom row) is shown. (A, C) Spatial distribution of channel sensitivity to the target ROI across tested scalp locations, color-coded by the average sensitivity over orientations. The ROI is highlighted in yellow on the cortical surface. (B, D) Scatter plots comparing ROI-level sensitivities estimated by photon propagation modeling (PPM) and the proposed CNN-based method.

## 4. Discussion

In this study, we propose a CNN-based framework for estimating subject-specific PMDFs from MR images and channel configurations. The proposed approach first trains a neural network to predict photon fluence distributions based on MR image and optode locations, and subsequently derives PMDFs using the predicted fluence distributions via the adjoint formulation. The framework was evaluated for channels placed over motor and frontal scalp regions across several commonly used source–detector distances. The results demonstrate high prediction accuracy across both cortical regions and channel configurations.

Compared with conventional photon propagation modeling methods, the proposed framework offers substantial advantages in computational efficiency. Monte Carlo–based approaches, while considered as the golden standard for PMDF estimation, require extensive computational resources due to the need for large numbers of photon simulations and accurate tissue segmentation. In contrast, the proposed method bypasses both explicit tissue segmentation and repeated photon simulations by directly learning the mapping from MR images to photon fluence distributions. Once trained, the model enables rapid PMDF estimation with significantly reduced computational cost. This capability is particularly important for real-time or subject-specific fNIRS applications, where rapid and accurate PMDF estimation is required.

Analytical models generally exhibit higher prediction errors due to their simplified assumptions, such as representing the head as a semi-infinite homogeneous medium. While analytical solutions are computationally efficient, their assumptions limit their ability to accurately capture the complex tissue structures of the human head. However, since the proposed method learned from the Monte Carlo simulation, its prediction naturally incoporate the complex tissue heterogeneous. Overall, the proposed approach achieves a balance between accuracy and efficiency, combining the realism of numerical methods with the speed of analytical models, thereby offering a practical solution for PMDF estimation in fNIRS applications.

The advantages of fast PMDF esimtion of the proposed framework also extend to various real-time fNIRS applications. For example, it enables interactive optode arrangement guided by PMDFs. Although algorithmic approaches for optode placement have been proposed, practical constraints in fNIRS systems—such as fixed source–detector distances and limited probe flexibility—often make it difficult to achieve optimal configurations prior to probe setup. In such cases, the ability to rapidly estimate PMDFs allows users to iteratively adjust optode positions with direct feedback on spatial sensitivity, without the need for repeated Monte Carlo simulations. In addition, fast PMDF estimation may benefit fNIRS-based neurofeedback, where accurate mapping between measured signals and underlying cortical regions is essential. By providing subject-specific spatial sensitivity information in real time, the proposed method can support more precise and interpretable feedback of brain activity.

Several limitations of the proposed framework should be acknowledged. First, the model was trained and evaluated on a subset of scalp locations and MR images drawn from the same population. As with many data-driven approaches, the performance of the model may degrade when applied to data that differ substantially from the training distribution. Although the results in the frontal region show only a slight reduction in accuracy— suggesting reasonable generalization within this population—further validation is required for broader applications. In particular, the performance of the model on MR images from different populations, such as subjects of varying ages or with pathological conditions, remains to be investigated. Second, the current model is trained for a single wavelength. However, optical properties of biological tissues are wavelength-dependent, and consequently the PMDF varies with wavelength. For example, longer wavelengths generally exhibit deeper tissue penetration and may result in higher PMDF values in cortical regions compared to shorter wavelengths. Extending the proposed framework to support multi-wavelength inputs would further enhance its applicability in practical fNIRS studies.

Future studies may focus on further improving the prediction accuracy of the proposed framework as well as expanding its range of applications. Recent work in related fields, such as deep learning–based estimation of electric field distributions in transcranial magnetic stimulation (TMS), has demonstrated that model performance can be enhanced through refining neural network architecture and training strategy (Li et al., 2022; Maki et al., 2025; Xu et al., 2021). Similar approaches could be explored to improve the accuracy and robustness of the proposed method. In addition, the framework can be extended to support multiple wavelengths and validated across more diverse populations, including infants and elderly subjects. Such extensions would improve its applicability in a wider range of fNIRS studies. Finally, the proposed method should be further evaluated in practical applications, including optode arrangement optimization, fNIRS data analysis, and image reconstruction in diffuse optical tomography (DOT).

## 5. Conclusion

In this study, we proposed a CNN-based framework for efficient estimation of photon measurement density functions (PMDFs) from subject-specific MR images and channel configurations. By learning to predict photon fluence distributions and deriving PMDFs via the adjoint formulation, the proposed method enable predict PMDFs of channels with flexible configurations without explicit tissue segmentation and computationally intensive photon propagation simulations. The results demonstrate that the framework achieves high accuracy in both voxel-level and ROI-level across multiple cortical regions and source–detector distances. Overall, this work presents an effective and scalable solution for PMDF estimation, with potential applications in optimizing optode arrangement, data analysis, and diffuse optical tomography.

## Conflict of Interest Statement

The authors declare no conflict of interest.

## Author contributions

**Yang Zhao**: Conceptualization, Methodology, Data Curation, Formal analysis, Writing -Original Draft. **Xue-Ting Sun**: Data Curation, Resources. **Wen-Di Shi**: Data Curation. **Chao-Zhe Zhu**: Conceptualization. **Lei Zhang**: Conceptualization, Supervision, Review & Editing.

## Data and code availability statements

The data and code that support the findings in this study can be available by contacting the corresponding authors, Prof. Lei Zhang, upon a formal data sharing agreement.

## Notes

### Competing Interest Statement

The authors have declared no competing interest.

### Funding

- Fundamental and Interdisciplinary Disciplines Breakthrough Plan of the Ministry of Education of China: JYB2025XDXM504

