## Supplementary Material for "Rapid estimation of photon measurement density functions using a deep convolutional neural network for functional near-infrared spectroscopy"

### 6. Supplementary Materials

#### 6.1. Validation of the MRI subvolume size

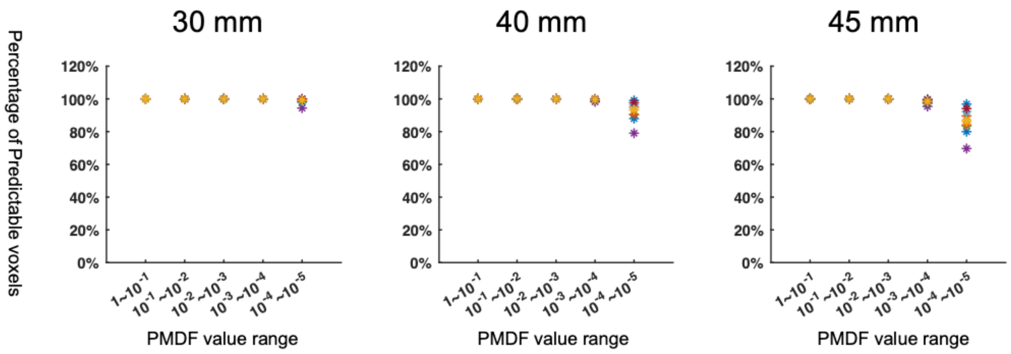


Figure S1. The percentage of predictable voxels across different PMDF value ranges for source-detector distances of 30, 40 and 45 mm.

We calculate the percentage of predictable voxels using a binary mask subvolume (all voxel values set to 1) and the PMDF derived from the PPM at 10-20 scalp locations. The predictable region for each channel was obtained by reslicing the mask subvolume into the native MRI space using the optode-centered coordinates of source and detector. The percentage of the predictable voxels within a given PMDF value range was computed as:

$$\frac{N_v}{N} \times 100\%, \quad v \in P,$$

where $N_v$ denotes the number of predictable voxels and $N$ is the total number of total voxels. For the 104 x 104 x 48 cuboid, the percentage of predictable voxels reaches 100% for PMDF values greater than $10^{-3}$ when source-detector distance is up to 45 mm. For commonly used 30 mm distance, voxels with PMDF values greater than $10^{-4}$ can be fully predicted. These results indicate that a 104 x 104 x 48 subvolume is sufficiently large for typical fNIRS channel configurations.
